# N-terminal region of *Drosophila melanogaster* Argonaute2 forms amyloid-like aggregates

**DOI:** 10.1101/2022.08.05.502916

**Authors:** Haruka Narita, Tomohiro Shima, Ryo Iizuka, Sotaro Uemura

## Abstract

Argonaute proteins play a central role in RNA silencing by forming protein-small RNA complexes responsible for the silencing process. While most Argonaute proteins have a short N-terminal region, Argonaute2 in *Drosophila melanogaster* (DmAgo2) harbors a long and unique N-terminal region. Previous *in vitro* biochemical studies have shown that the loss of this region does not impair the RNA silencing activity of the complex. However, an N-terminal mutant of *Drosophila melanogaster* has demonstrated abnormal RNA silencing activity. To explore the causes of this discrepancy between *in vitro* and *in vivo* studies, we investigated the biophysical properties of the region. Because the N-terminal region is highly rich in glutamine and glycine residues, which is a well-known property for prion-like domains (PrLD), the possibility of the N-terminal region functioning as a PrLD was tested. Our biochemical assays demonstrated that the N-terminal region can form aggregates that are not dissociated even in the presence of SDS. Also, the aggregates enhanced the fluorescence intensity of thioflavin-T, an amyloid detection reagent. The kinetics of the aggregation followed that of typical amyloid formation exhibiting the self-propagating activity. Further, we directly visualized the aggregation process of the N-terminal region under fluorescence microscopy and found that the aggregations took fractal or fibril shapes. Together, the results indicate that the N-terminal region is a PrLD. Many other PrLDs have been reported to modulate the function of proteins through their aggregation. Therefore, our results raise the possibility that aggregation of the N-terminal region regulates the RNA silencing activity of DmAgo2.

## Introduction

RNA silencing is involved in various biological processes including antiviral defense, development, and the maintenance of genomic integrity (1). Among the RNA silencing machinery, Argonaute proteins play key roles by directly binding to small RNAs to form the RNA Induced Silencing Complex (RISC) (2). After the small RNA in RISC recognizes its RNA targets, Argonaute proteins display endonucleolytic activity or translational repression (3). Despite the functional variety of Argonaute proteins, their structural organization is highly conserved, sharing four distinct domains: N, PAZ, MID, and PIWI (4). Previous studies have revealed the functions of each domain. The N domain initiates duplex unwinding immediately after small RNA binds to Argonaute (5). The PAZ and MID domains recognize the 5′ and 3′ ends of the small RNA, respectively (6–8). The PIWI domain, which shares structural similarities with ribonucleases, cleaves the target RNA(9, 10).

In contrast to these highly conserved domains, the N-terminal region, which is located upstream of the N domain, shows extensive diversity among species (11). While most Argonaute proteins have a short N-terminal region (e.g., 24 residues in human Ago2), Argonaute2 in *Drosophila melanogaster* (DmAgo2) harbors a long and unique N-terminal region. The N-terminal region (residues 1–398; henceforth, Nter) comprises almost one-third of the residues in DmAgo2 (1208 residues). Partial truncation of Nter (residues 326–371 deletion) has been reported to impair RNA silencing in mutant flies (12). However, an *in vitro* study demonstrated that another DmAgo2 mutant with Nter deletion (residues 1–278 deletion) still forms RISC and retains RNA cleavage activity(13).

Many eukaryotic RNA-binding proteins contain a prion-like domain (PrLD), which organizes intracellular condensates and regulates biochemical reactions (14–17). PrLDs consist of intrinsically disordered, low-complexity sequences often enriched in glutamine and glycine residues (18, 19). The Nter sequence shares this property. The content of glutamine and glycine residues in DmAgo2 Nter reach approximately 40% and 20%, respectively (11, 12). Inspired by these findings, here we tested whether DmAgo2 Nter is a PrLD. Our *in silico* prediction and biochemical assays demonstrated that Nter is a PrLD that can form amyloid-like aggregates. We also directly visualized by fluorescence microscopy how Nter aggregates and found that the aggregates are polymorphic. Since many PrLDs have been shown to regulate biochemical reactions through their aggregate formation (14–17), our results raise the possibility that Nter regulates RNA silencing activities in cells by forming aggregates.

## Results

### PrLD prediction based on the amino acid sequence of DmAgo2

To confirm the characteristics of DmAgo2 based on its amino acid sequence (Figure 1A), we performed intrinsically disordered regions (IDRs) prediction and three kinds of PrLDs prediction. First, to confirm that DmAgo2 harbors IDRs, we analyzed the DmAgo2 amino acid sequence using PONDR (20), a neural network-based prediction algorithm of IDRs. Two parts of Nter (residues 1−87 and 105−412) were identified as regions with high disorder scores (Figure 1B). Then, the possibility that Nter is not only an IDR but also a PrLD that can form amyloid-like aggregates was predicted by three PrLD prediction algorithms: the prion-like amino acid composition (PLAAC) (21), Prion Aggregation Prediction Algorithm (PAPA) (22), and PrionW (23). PLAAC, a Hidden Markov Model (HMM)-based prion prediction algorithm, identified residues 14−386 in Nter as the region with a high prion-like score (Figure 1B). Because the predictions of Nter being PrLD varied across the algorithms (Supplementary Figure 1), we next tested experimentally whether Nter has the capacity to form amyloid-like aggregates.

**Figure 1.**
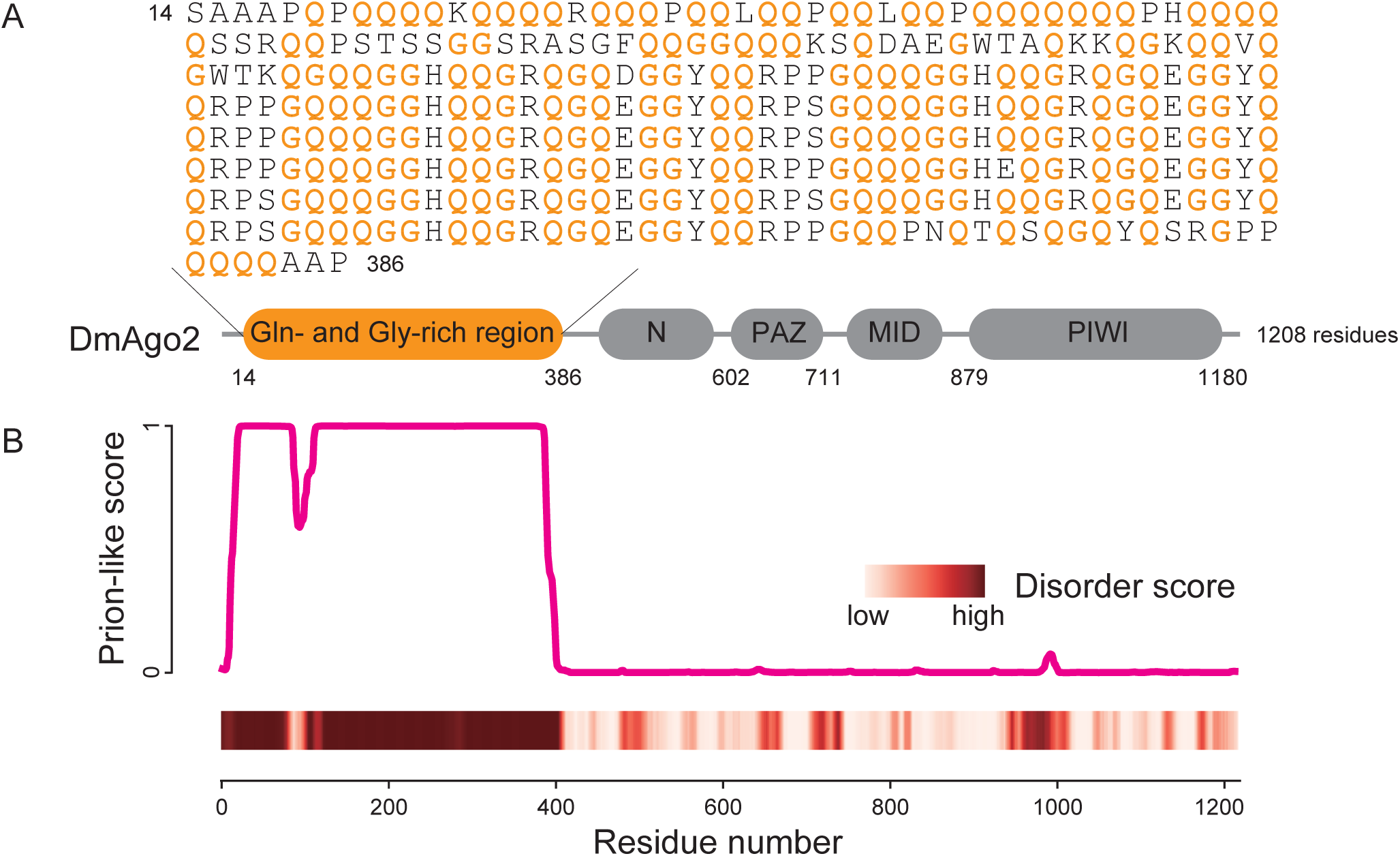
DmAgo2 has a long characteristic N-terminal region. **(A)** A schematic diagram of the full-length DmAgo2 protein and the sequence of N-ter. The four conserved domains are shown in gray, while the N-terminal region is shown in orange. In the N-ter sequence, glutamine and glycine residues are shown in orange. **(B)** Plot of the prion-like probability of the DmAgo2 sequence predicted by PLAAC (top) and a heatmap of the degree of disorder predicted by PONDR analysis (bottom).

### Nter can form SDS-resistant amyloid-like aggregates in a typical amyloid formation manner

We prepared a recombinant protein of Nter fused with mCherry at its N-terminal (mCherry-Nter) to test whether Nter behaves as a PrLD. PrLDs are known to form aggregates even in the presence of SDS (24). To test if Nter forms SDS-resistant aggregates, we first incubated 5 μM of monomeric mCherry-Nter in aggregation buffer (20 mM sodium phosphate, 50 mM sodium chloride, pH 7.4) for 3 days at room temperature. Then, the aggregates in the solution were collected by ultracentrifugation and subjected to semi-denaturing detergent agarose gel electrophoresis (SDD-AGE) (24). As references for the assay, mCherry and Sup35NM-mCherry were also subjected to SDD-AGE. mCherry is a monomeric protein (25), whereas Sup35NM is a well-known PrLD and can form amyloid-like aggregates (26). mCherry did not appear as a distinct band, indicating that mCherry did not form any aggregates (Figure 2). Contrarily, Sup35NM-mCherry and mCherry-Nter predominantly migrated as smears with high molecular weights, suggesting that mCherry-Nter formed SDS-resistant aggregates the same as Sup35NM-mCherry (Figure 2). The aggregates of Sup35NM-mCherry and mCherry-Nter were dissolved into soluble fractions when boiled. Thus, the results showed that Nter forms SDS-resistant aggregates.

**Figure 2.**
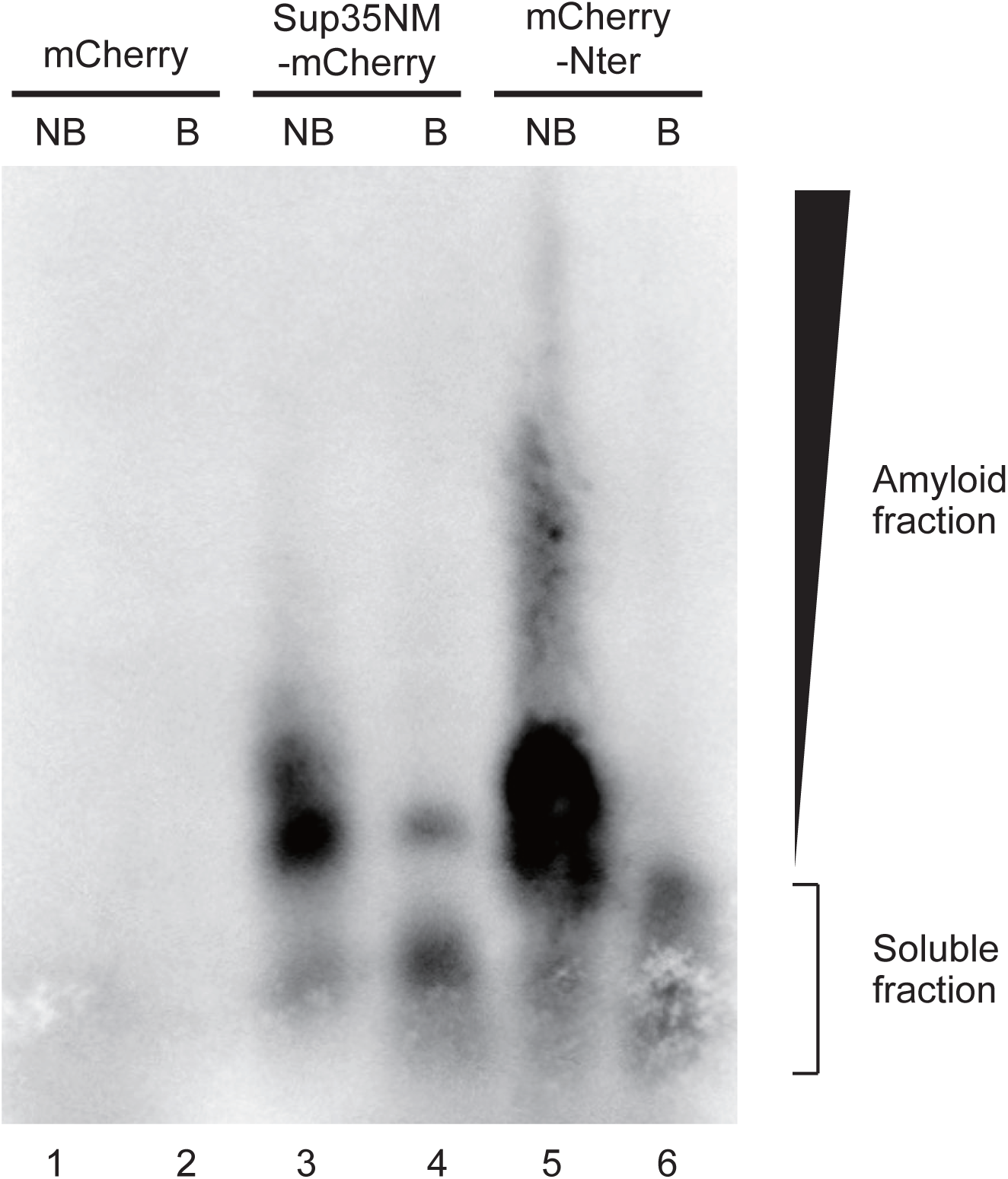
mCherry-Nter formed SDS-resistant aggregates. SDD-AGE analysis of mCherry-Nter aggregates. mCherry and Sup35NM-mCherry were used as reference controls. mCherry-tagged polypeptides were detected by western blotting with anti-RFP polyclonal antibodies. NB, non-boiled fraction; B, boiled fraction.

To further test whether Nter aggregates take amyloid-like structures, we performed Thioflavin-T (ThT) assays. ThT is a fluorescent probe with specific fluorescence enhancement upon binding with amyloid fibrils (27, 28). First, we measured the fluorescence spectrum of 20 μM ThT with and without 5 μM Nter solution incubated for 12 h beforehand. In the presence of Nter, the fluorescence intensity of ThT was enhanced approximately four times compared to that without Nter, and a peak at 490 nm was clearly visible in the fluorescence spectrum (Figure 3A), indicating the presence of amyloid-like structures in the Nter solution.

**Figure 3.**
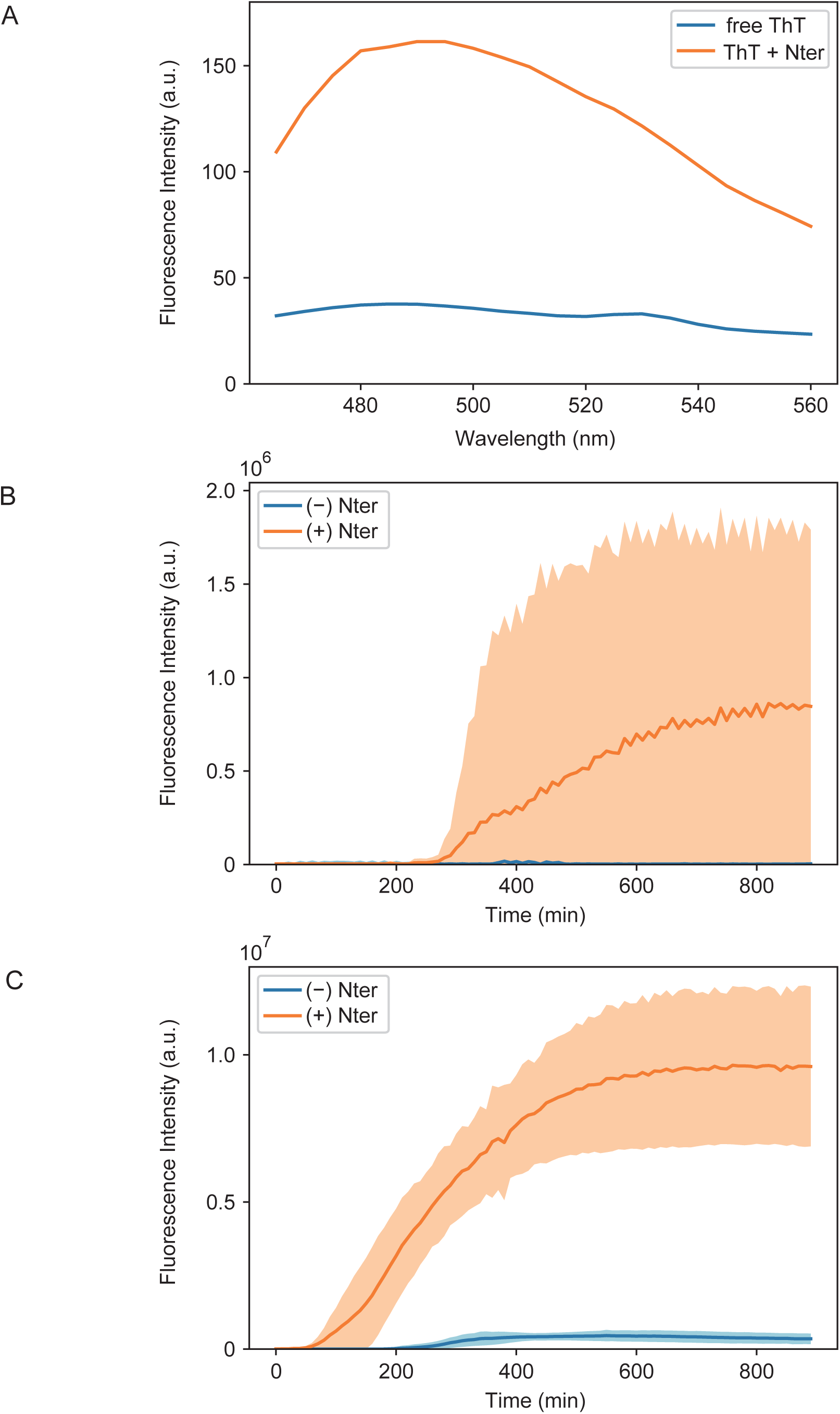
Nter aggregated in the same manner as common amyloids. (**A**) Fluorescence spectra of ThT in the absence (blue) and presence (orange) of Nter aggregates. (**B**) Time-course of the ThT fluorescence intensity in the absence (blue) and presence (orange) of mCherry-Nter monomers. The solid curves and filled area indicate the mean value and standard deviation of three repetitive experiments, respectively. (**C**) Time-course of ThT fluorescence intensity in the absence (blue) and presence (orange) of mCherry-Nter seeds. The solid curves and filled area indicate the mean value and standard deviation of three repetitive experiments, respectively.

To confirm that these aggregates were formed in a typical amyloid formation manner, we tracked the aggregation process over time by monitoring the ThT fluorescence intensity under microscopy. Consistent with the canonical amyloid formation from monomers generally requiring a lag phase for nucleation (29), the ThT fluorescence intensity with Nter showed an initial lag phase of ~300 min and then reached a plateau at ~800 min after mixing monomeric Nter with ThT in aggregation buffer (Figure 3B, Supplementary Figure 2A). Prion-like proteins generally exhibit seeded self-propagation *in vivo* and *in vitro* (26, 30). Therefore, we prepared the seeds of Nter aggregates and examined whether the seeds can accelerate the aggregation process. The addition of the seeds reduced the initial lag phase to less than 100 min (Figure 3C, Supplementary Figure 2B), showing that the Nter aggregation process follows canonical amyloid formation kinetics with a self-propagation property.

### Direct fluorescence visualization of Nter aggregates

Next, we directly visualized the shape of Nter aggregates using fluorescence microscopy to investigate their morphology. The fluorescence of mCherry-Nter that had been flushed into a glass chamber and incubated for 12 h to form aggregates revealed fractal or fibril-shaped aggregates. When ThT was added to the chamber, the fluorescent images of ThT and mCherry matched completely, suggesting that the entire region of the Nter aggregates took an amyloid-like structure (Figure 4A−D).

**Figure 4.**
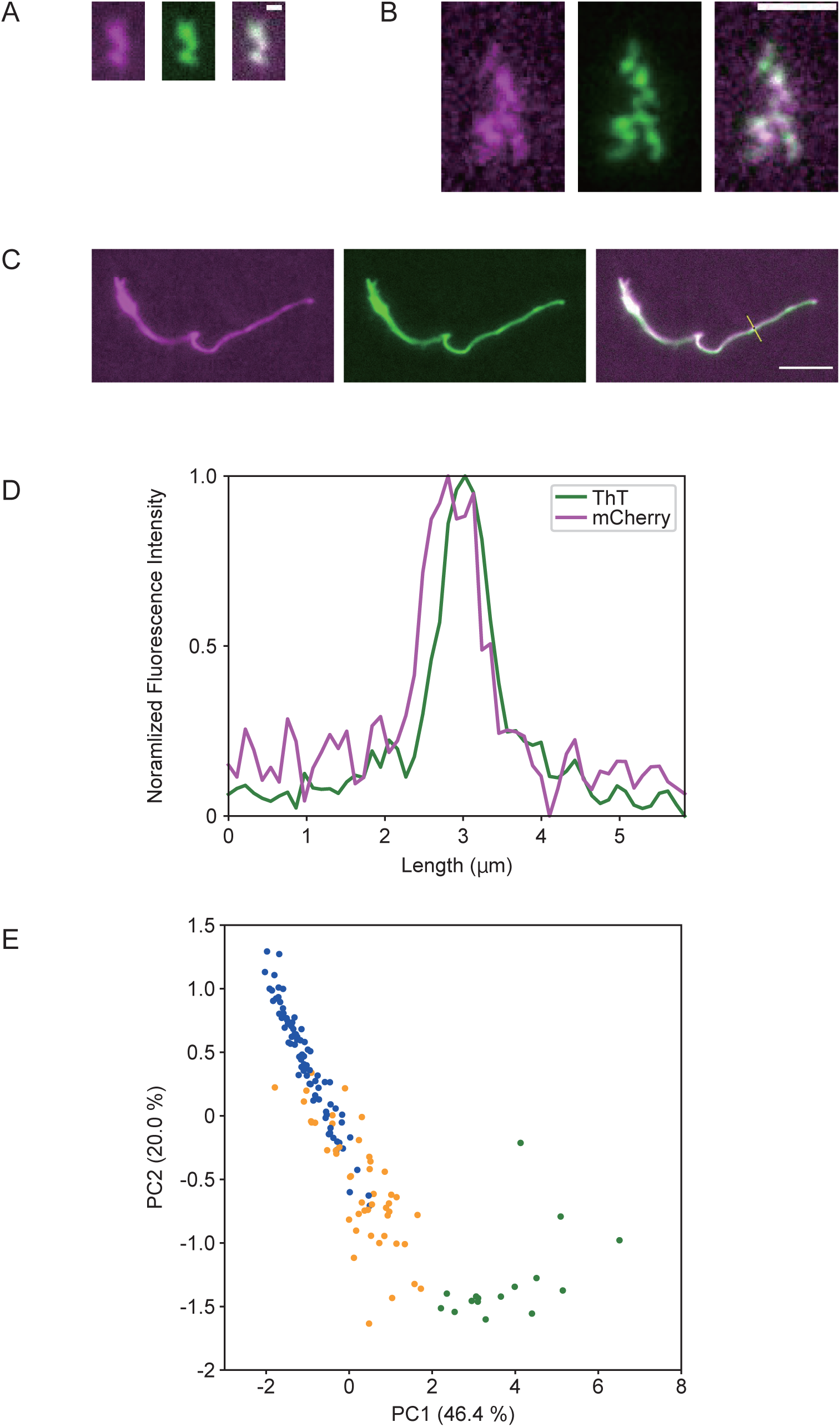
Fluorescence imaging of the Nter aggregate morphology. (**A**–**C**) Representative fluorescent images of linear fractal-shaped (**A**), branched fractal-shaped (**B**) and fibril-shaped aggregates (**C**). Fluorescence signals from mCherry (left panels, magenta) and ThT (middle panels, green) showed almost identical images. Merged images are shown in the right panels. Scale bars represent 1 μm (**A**, **B**) and 5 μm (**C**). (**D**) Fluorescence intensity profiles of mCherry and ThT along the line in the merged image of Figure 4C. Intensities were normalized. **(E)** PCA analysis of the shapes of the Nter aggregates. The plots were colored according to the classified group (blue, linear fractal-shaped; green, branched fractal-shaped; orange, fibril-shaped).

Since the aggregates showed various shapes (Figure 4A–C and Supplementary Figure 3, we analyzed and classified the shapes using principal component analysis (PCA) with k-means clustering into three groups: linear fractal-shaped, branched fractal-shaped and fibril-shaped aggregates (Figure 4E). The most frequently observed group (56%) was linear fractal-shaped aggregates (Figure 4A and Supplementary Figure 3A). In these aggregates, ~0.5 μm fluorescent puncta were linearly connected to each other. The median length of these aggregates was 1.4 [1.2–2.1] μm (median with quartile range, *n* = 83, Supplementary Figure 4A), corresponding to ~2 puncta. For the second-largest group, 32% of the whole aggregates were classified as branched fractal-shaped aggregates (Figure 4B and Supplementary Figure 3B). In the aggregates of this group, several puncta were bound to branch off from the main chain of the aggregates. The median length of the longest chain of the aggregates was 3.3 [2.2–4.9] μm (median with quartile range, *n* = 47, Supplementary Figure 4B), and the median number of branched points was 4 [3–6] (median with quartile range, *n* = 47, Supplementary Figure 4C). The remaining 12% of the whole aggregates were classified as fibril-shaped aggregates (Figure 4C and Supplementary Figure 3C). This fibril shape is the most common structure for amyloids (31). The median length of the Nter fibril was 27 [19–50] μm (median with quartile range, *n* = 17, Supplementary Figure 4D), taking a shape much longer than the fractal-shaped aggregates. Most Nter fibrils contained a few number of puncta, but these puncta did not directly bind to each other. Thus, we could classify the shapes of the Nter aggregates according to the presence of branching, the length of the aggregates and the alignment of puncta.

Time-lapse imaging of the Nter aggregation process demonstrated how the fractal-shaped aggregates formed. First, fluorescent puncta docked to each other, forming linear fractal-shaped aggregates (Figure 5A). Next, linear fractal-shaped aggregates docked to each other, forming branched fractal-shaped aggregates (Figure 5B). Finally, branched fractal-shaped aggregates docked to each other, forming larger branched fractal-shaped aggregates (Figure 5C). Therefore, it is most likely that the linear and branched fractal-shaped aggregates share the same formation process, and the difference in the stage of the formation process is seen as the difference in shape.

**Figure 5.**
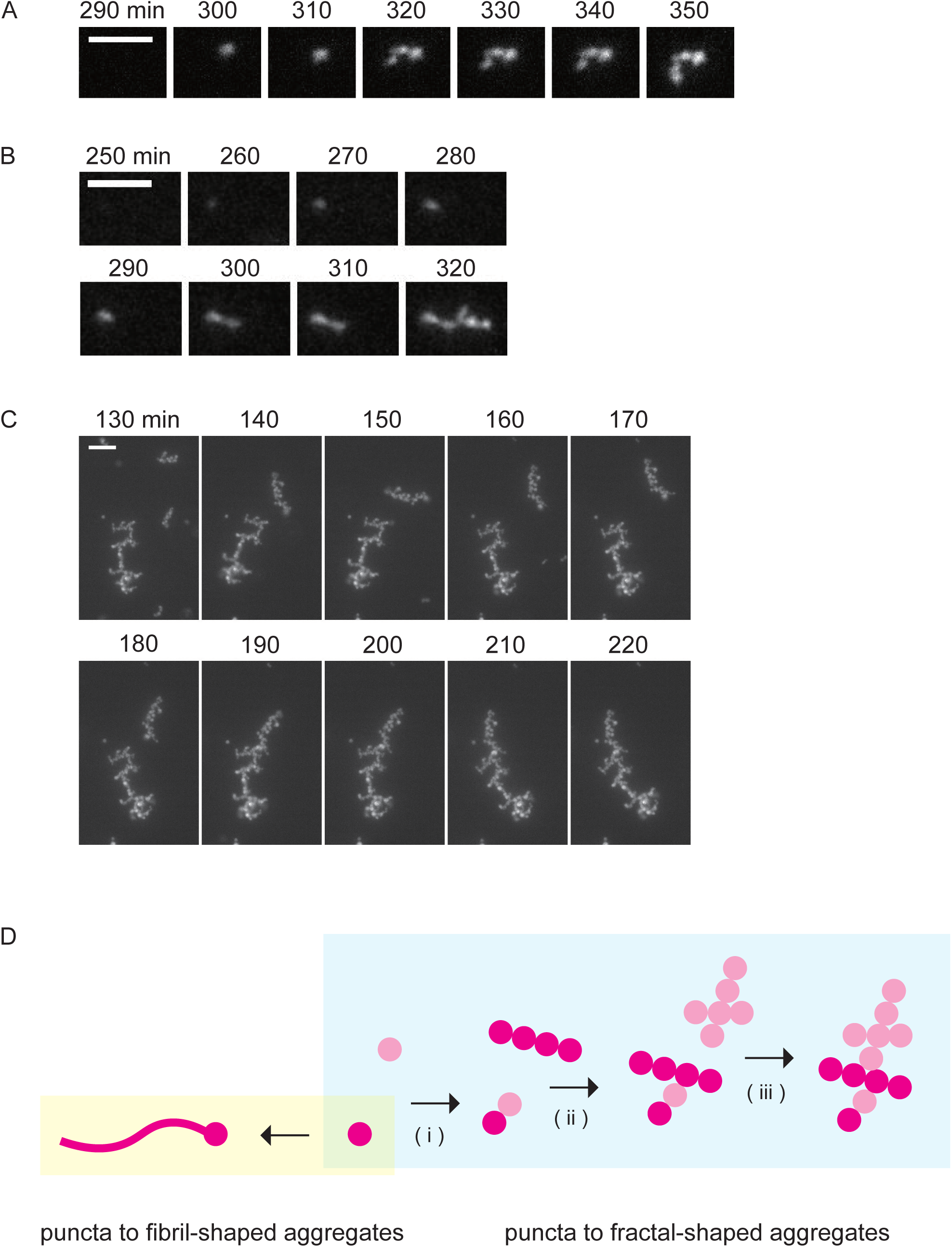
Fractal-shaped Nter aggregates grew by docking between puncta. (**A–C**) Time-lapse images of the Nter aggregates formation. Ten-minute intervals are shown from left to right. Scale bars represent 5 μm (**A, B**) and 10 μm (**C**). (**D**) Model of the aggregate formation. Assuming both fibril- and fractal-shaped Nter aggregates grew from a single punctum, the polymerization of Nter monomers from the punctum results in fibril-shaped aggregates, while the fractal-shaped aggregates result from puncta binding to each other (i). The binding of several linear fractal-shaped aggregates produces branched fractal-shaped aggregates (ii). The branched fractal-shaped aggregate becomes bigger by docking with other fractal-shaped aggregates (iii).

## Discussion

Here, we demonstrated that Nter is a PrLD by *in silico* prediction and biochemical assays. The PLAAC algorithm predicted Nter is a PrLD based on its characteristic sequence. mCherry-Nter fusion protein formed aggregates that exhibit amyloid-specific properties of resistance to SDS and binding to ThT. Furthermore, Nter aggregated in a manner consistent with typical amyloids, including a lag phase for nucleation and self-propagation activity. These results strongly suggest that Nter can form amyloid-like aggregates.

From a structural viewpoint, we found that Nter aggregates can take two distinct conformational states. Most aggregates took fractal shapes, while the remaining had fibril shapes. This observation is consistent with previous studies showing that fractal-shaped aggregates tend to form under diffusion limited-conditions (32–35), because protein diffusion is limited in the 145-μm thick chamber in our assays. Within cells, protein diffusion should be even more restricted due to molecular crowding and the size of the cytoplasm. Therefore, Nter aggregates are expected to be more prone to taking on a fractal shape, although it is yet to be demonstrated that these observed fractal-shaped aggregates are really formed in cells. In addition, fractal-shaped aggregates docked to form larger fractal-shaped aggregates. On the other hand, although the forming process of fibril-shaped aggregates was not observed, fibril-shaped aggregates frequently contained a few numbers of puncta in their fibrils (Figure 4C and Supplementary Figure 3C). This finding implies that fibril-shaped aggregates can elongate from the edges of the puncta. Thus, we propose a growth model of Nter aggregates in which the docking of several puncta form fractal-shaped aggregates and polymerization of monomeric Nter from a single punctum forms fibril-shaped aggregates (Figure 5D).

Nter aggregation may indirectly modulate the RNA silencing activity of DmAgo2. It has been shown that Nter is not essential for DmAgo2 to exhibit RNA silencing activity under *in vitro* conditions, but the partial truncation of Nter causes RNA silencing defects in the compound eyes of flies (12). Combining these previous studies with our current results raises the possibility that *in vivo* Nter aggregation modulates RISC formation and/or RNA cleavage activity by other domains of DmAgo2. Protein aggregation can both be inhibitory on protein activity by steric hindrance effects on the substrate binding or facilitative by increasing local protein concentrations (36). Furthermore, it has been reported that protein aggregation is a major driving force for liquid-liquid phase separation in the cytoplasm, allowing for the proper subcellular localization of biomolecules (14). Indeed, DmAgo2 is localized in cytoplasmic D2 bodies, where endogenous siRNAs are loaded onto DmAgo2, and this localization is essential for the formation of RISC with the proper small RNA (37). The ability of Nter to form amyloid-like aggregates may contribute to such modulation of DmAgo2 activity and localization. Although it still needs to be verified whether the aggregation of endogenous DmAgo2 actually occurs in cells, this hypothesis can explain the apparent inconsistency among previous *in vitro* and *in vivo* studies regarding the effects of Nter in RNA silencing.

We focused on DmAgo2 because *Drosophila melanogaster* is one of the most well-studied model organisms in RNA silencing studies. However, the aggregation of the N-terminal region may be a general phenomenon not limited to DmAgo2. Although Argonaute proteins of many organisms have only a short N-terminal region, Ago2 among arthropods share glutamine- and glycine-enriched properties in the N-terminal region (38). Their amino acid composition, which is a major factor in amyloid-formation potential (39), is close to DmAgo2 Nter. Therefore, the N-terminal regions of Ago2 in other arthropods may also form amyloid-like aggregates as PrLDs. Also, some plant Argonaute proteins possess glutamine- and glycine-rich N-terminal regions (Supplementary Figure 5), implying they share the same property. Since many Argonaute proteins are likely to have the similar characteristic sequence at the N-terminus, we propose to call such region N-terminal Argonaute Prion-like (NAP) domain. The physiological features and function of this aggregation property of NAP domains await further study, but the property has the potential to function as a regulatory mechanism for RNA silencing across species.

## Experimental procedures

### In silico analysis

The amino acid sequence of DmAgo2 (equivalent to Uniprot entry Q9VUQ5 with the deletion of residues 43–48) was derived from native *Drosophila* DNA. The sequence was analyzed using the programs PLAAC (http://plaac.wi.mit.edu/) (21), PAPA (https://combi.cs.colostate.edu/supplements/papa/) (40), and PrionW (http://bioinf.uab.cat/prionw/) (23) with default settings. Disorder scores were calculated using the Predictor of Natural Disordered Regions (PONDR) algorithm with the VLXT predictor (http://www.pondr.com/) (41).

### Plasmid construction

The expression plasmid for N-terminal 6×His-tagged mCherry-Nter was constructed by inserting the sequence encoding DmAgo2 Nter (residues 1–398) amplified from the native DmAgo2 sequence into pCold I (TAKARA Bio) using the InFusion HD cloning kit (Clontech). The expression plasmid for C-terminal 6×His-tagged Sup35NM-mCherry was constructed by inserting the sequence encoding mCherry into the expression plasmid for Sup35NM based on pET29b (residues 1–253) (a kind gift from Prof. M. Tanaka (RIKEN CBS, Japan)).

### Protein expression and purification

For the protein expression, *E. coli* BL21(DE3) cells were transformed with the constructed plasmids and selected using 50 mg/L carbenicillin (for pCold I and pET11a plasmids) or 50 mg/L kanamycin (for pET29b plasmid). These antibiotics were added in all culture media described below. The transformed *E. coli* cells were pre-cultured in 5 mL of LB medium overnight at 37°C and then inoculated into 2 L of TB medium. The cells were grown at 37°C until OD_600_ reached 1.2 (6×His-tagged mCherry-Nter and 6×His-tagged Sup35NM-mCherry) or 0.6 (mCherry-His). 6×His-tagged mCherry-Nter was expressed with 1 mM IPTG at 12°C. mCherry-His and 6×His-tagged Sup35NM-mCherry were expressed with 1 mM IPTG at 28°C. After 20 h of culture, the cells were harvested and then lysed in 80 mL of lysis buffer (20 mM Hepes-KOH pH8.0, 100 mM NaCl, and 6 M GdnHCl) with 10-s sonication at an intensity of 2 on ice using the ultrasonic disruptor (UD211, TOMY) a total of 9 times with 20 s intervals in between. Following centrifugation of the lysate at 18,000 × g for 30 min, 6 mL of 2 × Ni-NTA resin (Ni-NTA Agarose HP, FUJIFILM Wako Pure Chemical) was added to the supernatant. After a 60-min rotation at room temperature, the solution was transferred to a disposal column (Muromac Mini-column M, Muromachi Chemical Inc). The resin was washed twice with 3 column volumes (CVs) of wash buffer (40 mM imidazole-HCl, 20 mM Hepes-KOH, pH8.0, 100 mM NaCl, and 6 M GdnHCl) and eluted with 1 CV of elution buffer (400 mM imidazole-HCl, 20 mM Hepes-KOH, pH8.0, 100 mM NaCl, and 6 M GdnHCl) three times. The elution fractions were resolved on 12% denaturing polyacrylamide gels and visualized by Coomassie Brilliant blue (Nacalai Tesque) staining. Elution fractions containing proteins of interest were pooled and ultra-filtered to remove large aggregates using Vivaspin 6, 100,000 MWCO (Sartorius Stedim Biotech GmbH). Finally, the flow-through was concentrated using Vivaspin 6, 30,000 MWCO (Sartorius Stedim Biotech GmbH). The purified proteins were snap-frozen in liquid nitrogen and stored at −80°C until use.

Monomers of mCherry-Nter, mCherry, and Sup35NM-mCherry were prepared as follows. First, purified proteins were diluted 50 times by aggregation buffer (20 mM sodium phosphate, pH 7.4, 50 mM NaCl). Next, the protein solution was ultracentrifuged at 418,000 × g for 20 min at 25°C to remove large aggregates. Following the ultracentrifugation, the supernatant, equivalent to a half volume of the solution, was immediately transferred to a new 1.5 mL tube. The protein concentrations were determined using the following molar extinction coefficients at 280 nm: mCherry-Nter, 66,240 M^−1^cm^−1^; Sup35NM-mCherry, 65,670 M^−1^cm^−1^; and mCherry, 35,870 M^−1^cm^−1^. The molar extinction coefficients were calculated using the ProtParam tool of ExPASy (https://web.expasy.org/protparam/) (42). Then, the protein solution was diluted to 5 μM by aggregation buffer. Nter aggregates were prepared by incubating the above solution at room temperature for the given time.

Seeds were prepared as follows. First, purified mCherry-Nter was diluted 10 times by elution buffer. Next, the solution was applied to a gel filtration spin column (Micro Bio-Spin 30 column, Bio-Rad) equilibrated with aggregation buffer to remove GdnHCl and ultracentrifuged at 418,000 × g for 20 min at 25°C to remove large aggregates. Finally, the supernatant, equivalent to 90% solution volume, was immediately transferred to a new 1.5 mL tube as a seed.

### SDD-AGE

After monomers were incubated for 3 days at room temperature without agitation, aggregates were pelleted by ultracentrifugation at 418,000 × g for 20 min at 25°C. After completely removing the supernatant, the pellet was resuspended with aggregation buffer. Prepared samples were subjected to SDD-AGE analysis. First, aggregate solutions were mixed with 4 × sample buffer (2 × TAE, 20% (v/v) glycerol, 4% (w/v) SDS, 0.25% (w/v) bromophenol blue). Next, after incubation for 15 min at room temperature with or without heat treatment at 95°C for 2 min, the samples were loaded onto a 1.5% agarose gel containing 1 × TAE and 0.1% SDS. Finally, the aggregates and monomer protein were resolved on the gel in running buffer (1 × TAE, and 0.1% SDS) at 75 V for 90 min at 4°C, followed by capillary blotting onto a PVDF membrane (FUJIFILM Wako Pure Chemical) for the western blotting analysis, as described previously (24). Anti-RFP polyclonal antibody (1/5,000 dilution from the product, PM005, MBL) and HRP-labeled IgG detector (1/5,000 dilution from the product, Western BLoT Rapid Detect v2.0, Takara Bio) were used as primary and secondary antibodies, respectively. Proteins were detected with SuperSignal West Femto (Thermo Scientific) on a gel imager (Amersham Imager 600, Cytiva).

### Measurement of ThT spectrum

5 μM mCherry-Nter monomers were incubated for 12 h at 25°C to prepare mCherry-Nter aggregates. Fluorescence spectra of mCherry-Nter aggregates in solution with or without 20 μM ThT (FUJIFILM Wako Pure Chemical) were obtained using a fluorescence spectrometer (RF-6000, Shimadzu) at room temperature. The excitation wavelength was set at 455 nm (bandwidth: 5 nm), and the emission was recorded from 465 to 560 nm (bandwidth: 5 nm).

### Chamber preparation for fluorescence microscopy

Flow chambers were prepared as described previously (Narita et al. 2020 *bioRxiv*) with some modifications. Briefly, the coverslips (No. 1S 22 × 22 mm and No.1S 24 × 32 mm, Matsunami) were cleaned in 1 N KOH for 15 min with sonication (Bransonic tabletop cleaner, Emerson). All subsequent preparation procedures were performed in a clean hood (Matsusada Precision). After 20 times rinsing with Milli-Q water and drying in a dryer, the coverslips were cleaned using a plasma cleaner (YHS-R, SAKIGAKE-Semiconductor Co., Ltd.). A 25 μL volume micro-chamber was made by placing a small coverslip of 22 × 22 mm over a 24 × 32 mm glass coverslip using double-sided adhesive tape (145 μm thickness, TERAOKA SEISAKUSHO CO., LTD) in a clean hood. First, Lipidure-BL103 (NOF Corporation) was flowed into the chamber to coat the glass surface. After a 2-min incubation and excess Lipidure-BL103 removal by three washes with 25 μL aggregation buffer, each sample was flushed into the glass chamber.

To evaluate the colocalization of ThT and mCherry and the size distribution of the aggregates, mCherry-Nter aggregates were prepared by a 12 h-incubation of monomer mCherry-Nter at room temperature. mCherry-Nter aggregates were observed as follows. mCherry-Nter aggregates solution was flushed into a Lipidure-coated glass chamber and incubated for 2 min at room temperature. Finally, excess mCherry-Nter unbound on the glass surface was removed by three washes with 25 μL aggregation buffer with 20 μM ThT.

To measure the aggregation kinetics with or without the aggregate seeds, 5 μM mCherry-Nter monomers were flushed into a Lipidure-coated glass chamber and sealed with VALAP (1:1:1 mixture of vaseline, lanolin, and paraffin). In the presence of seeds, the final 5 μM of seeds was added to the monomer solution.

### Fluorescence Microscopy

Nter aggregates were observed using an inverted microscope (Nikon Ti-E) equipped with a filter cube for mCherry imaging consisting of an excitation filter (#67-033, Edmund Optics), a dichroic mirror (#67-083, Edmund Optics) and an emission filter (#67-036, Edmund Optics) and a filter cube for ThT imaging consisting of an excitation filter (#67-026, Edmund Optics), a dichroic mirror (#67-078, Edmund Optics) and an emission filter (#67-028, Edmund Optics). mCherry and ThT were illuminated with a 532 nm-laser (OBIS LX/LS, Coherent) and 445 nm-LED light (SOLIS-445C, THORLABS), respectively. The images were obtained through a Plan Fluor 10×/0.30 or Plan Apoλ 40×/0.95 objective (Nikon) and recorded at 10 frames/s using an Orca Flash4.0 V3 digital CMOS camera (Hamamatsu Photonics). All equipment in the microscopy system was controlled by Micro-Manager software (43). To measure aggregates formation kinetics, ThT fluorescence images were recorded at 10-min intervals.

### Image Analysis

Image analysis was performed using either ImageJ or Python. To accurately evaluate the colocalization of ThT and mCherry, we corrected the chromatic aberration of the images obtained from both fluorescence channels using ImageJ and the TurboReg plugin (http://bigwww.epfl.ch/thevenaz/turboreg/). The translational distortion parameters were obtained by comparing the averaged image from 10 images of an objective micrometer (OB-M, 1/100, Olympus) in each channel. Four regions of interest (3 × 3 pixels) were used for the calculation to minimize artifacts. The images of mCherry-Nter aggregates corrected using the obtained parameters were used for further colocalization analysis.

To classify the shapes of the aggregates observed by the ThT fluorescence, we first extracted features listed below: the longest chain length, the mean width, the mean width per longest chain length, the total fluorescence intensity, the number of branched points per longest chain length, the number of fluorescence intensity peaks per longest chain length, and the percentage of the longest chain length to total length. To calculate these features, the images were preprocessed as described below. First, the background of each image was subtracted using the “Subtract Background” algorithm in ImageJ with a rolling ball radius of 200 pixels. Next, bright particles with more than 9 pixels with fluorescence intensity beyond the given threshold (60 a.u. out of 65,535 a.u.) were detected as aggregates. Following the above preprocessing, to calculate the total length of each aggregate, the “Skeletonize” algorithm was applied to each detected aggregate. Next, the longest chain length and the number of branched points of each aggregate were calculated from the skeletonized images. The mean width of each aggregate along the longest chain was calculated by averaging the widths along the longest chain using ImageJ. The number of fluorescence intensity peaks per longest chain length was detected using the Laplacian of Gaussian (LoG) method available in the scikit-image module in Python (44). This number was evaluated because the smallest unit of a fractal-shaped aggregate is a round-shaped aggregate detected as a fluorescence punctum. Finally, we performed a PCA analysis based on the extracted eight features to obtain the two axes corresponding to the maximum variation (PC1) and second-most variation (PC2). Using the plot, the k-means clustering algorithm classified these aggregates into four groups. One of these groups consisted of only one sample, where both PC1 and PC2 values were outliers because two ends of the aggregate were connected to form a ring. We re-classified this one sample into “fibril-shaped aggregate”. The PCA result was plotted using a Python plotting library (Matplotlib: http://matplotlib.org) (45).

To calculate the amount of ThT bound to the aggregates at each time point, bright particles with more than 9 pixels of high fluorescence intensity (more than 250 a.u. out of 65,535 a.u.) were detected as the aggregates. ThT fluorescence intensity of the pixels with the detected aggregates were plotted against time.

## Data availability

All image data sets and source codes for image analysis were deposited at Figshare (10.6084/m9.figshare.20438139)

## Acknowledgement

We thank Dr. Motomasa Tanaka (RIKEN Center for Brain Science) for providing the expression plasmid of Sup35NM. We also thank the members of the Uemura and Siomi laboratories for valuable discussions.

## Author contributions

Conceptualization, H.N. and T.S.; Data curation, H.N.; Methodology, H.N. and T.S.; Investigation, H.N.; Analysis, H.N.; Supervision, T.S.; Visualization, H.N. and T.S.; Writing – original draft, H.N. and T.S.; Writing – review & editing, H.N., T.S., R.I. and S.U.

## Funding and additional information

HN is supported by a JSPS Research Fellowship for Young Scientists and WINGS-LST, the University of Tokyo. This work is supported by Grant-in-Aid for JSPS Fellows, 21J11218 (to HN) and by JSPS KAKENHI (18K06147,19H05379 and 21H00387 to T.S.).

