## Supporting Information for "N-terminal region of *Drosophila melanogaster* Argonaute2 forms amyloid-like aggregates"

**Contents,**

**Supplementary Figure S1.** PAPA did not predict Nter as PrLD

**Supplementary Figure S2.** Time-lapse images of N-ter aggregated in the glass chamber

**Supplementary Figure S3.** Representative images of N-ter aggregates

**Supplementary Figure S4.** Characteristics of each type of N-ter aggregate

**Supplementary Figure S5.** Some plant Argonautes are also predicted to have a NAP domain

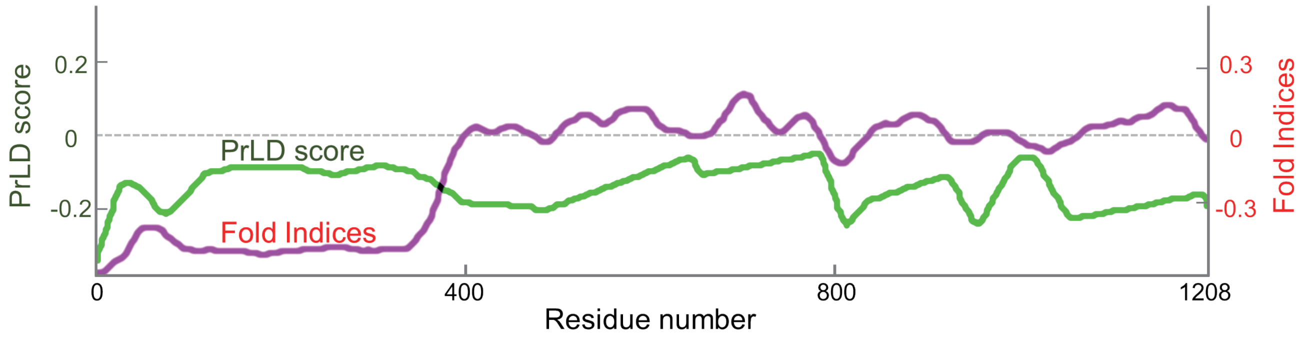


**Supplementary Figure S1. PAPA did not predict Nter as PrLD**

The expected PrLD scores of DmAgo2 Nter predicted by PAPA. The green line indicates prion formation propensity and the magenta line indicates fold indices.

A

B

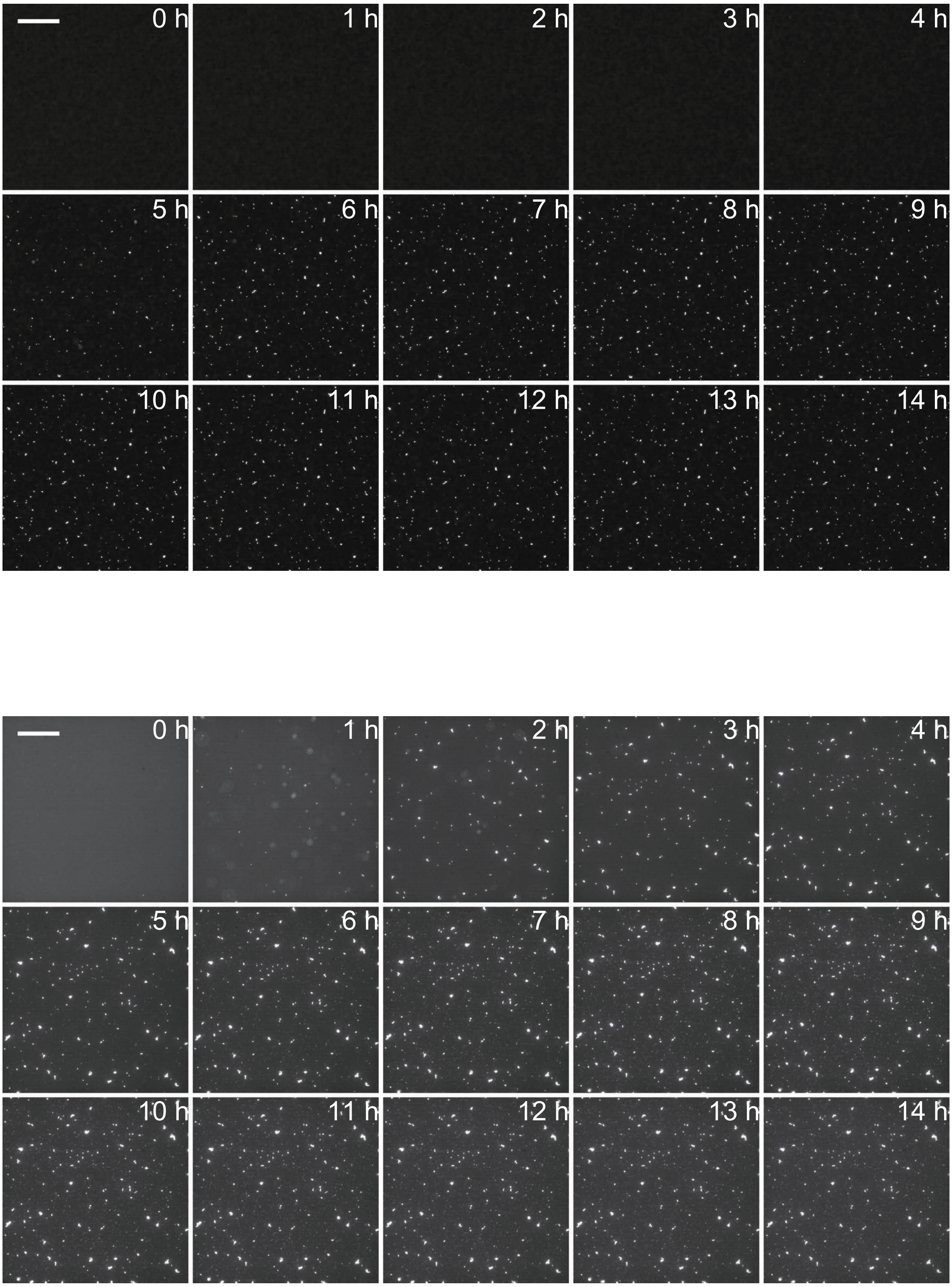


**Supplementary Figure S2. Time-lapse images of N-ter aggregated in the glass chamber**

Fluorescent images of ThT with 5 μM mCherry-Nter in the absence **(A)** and presence **(B)** of seeds. Time interval between images is 1 h. Scale bars, 50 μm.

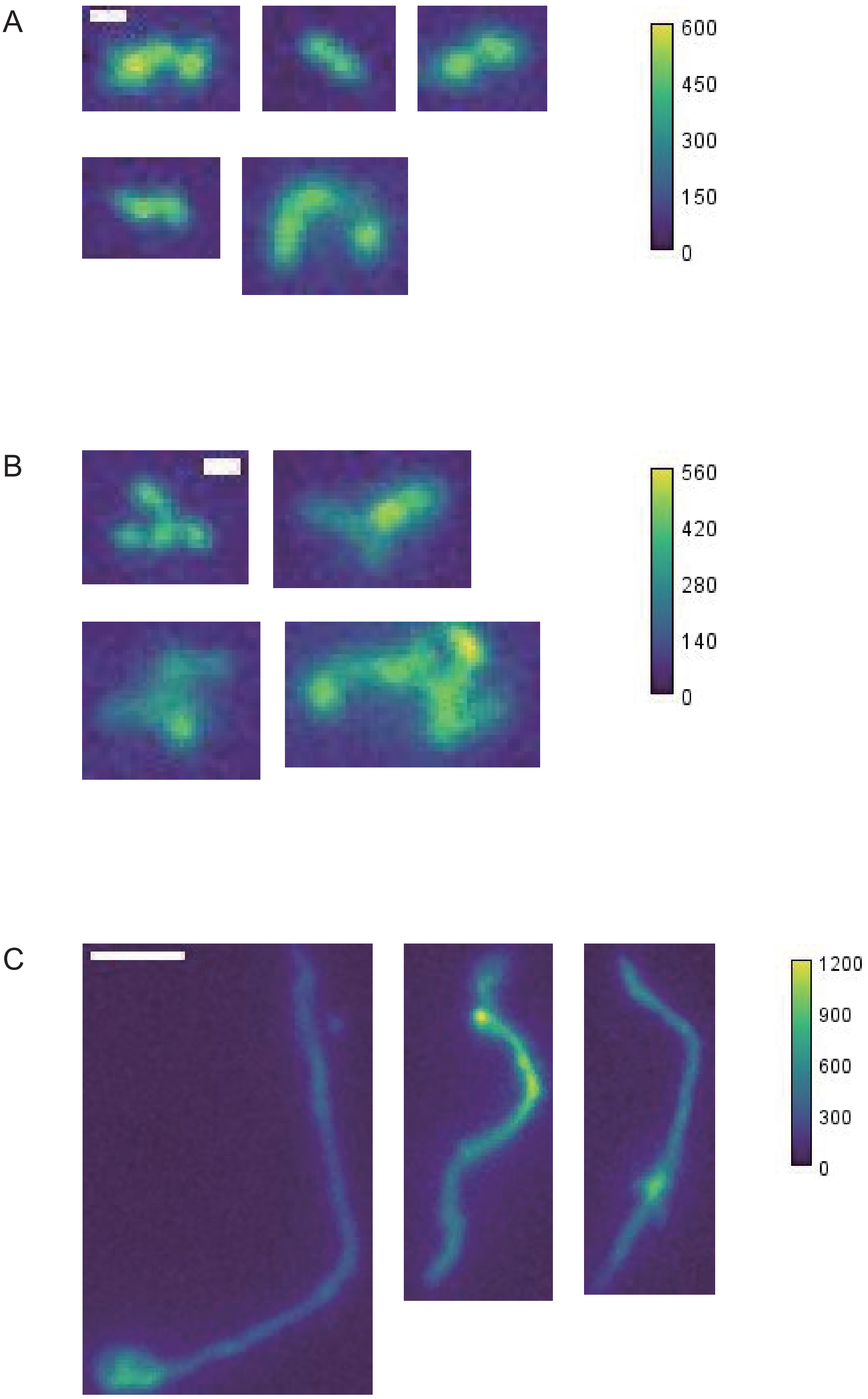


**Supplementary Figure S3. Representative images of N-ter aggregates**

ThT fluorescent images of linear fractal-shaped **(A)**, branched fractal-shaped **(B)** and fibril-shaped **(C)** N-ter aggregates. The color bar shows the fluorescence intensity (a.u.).

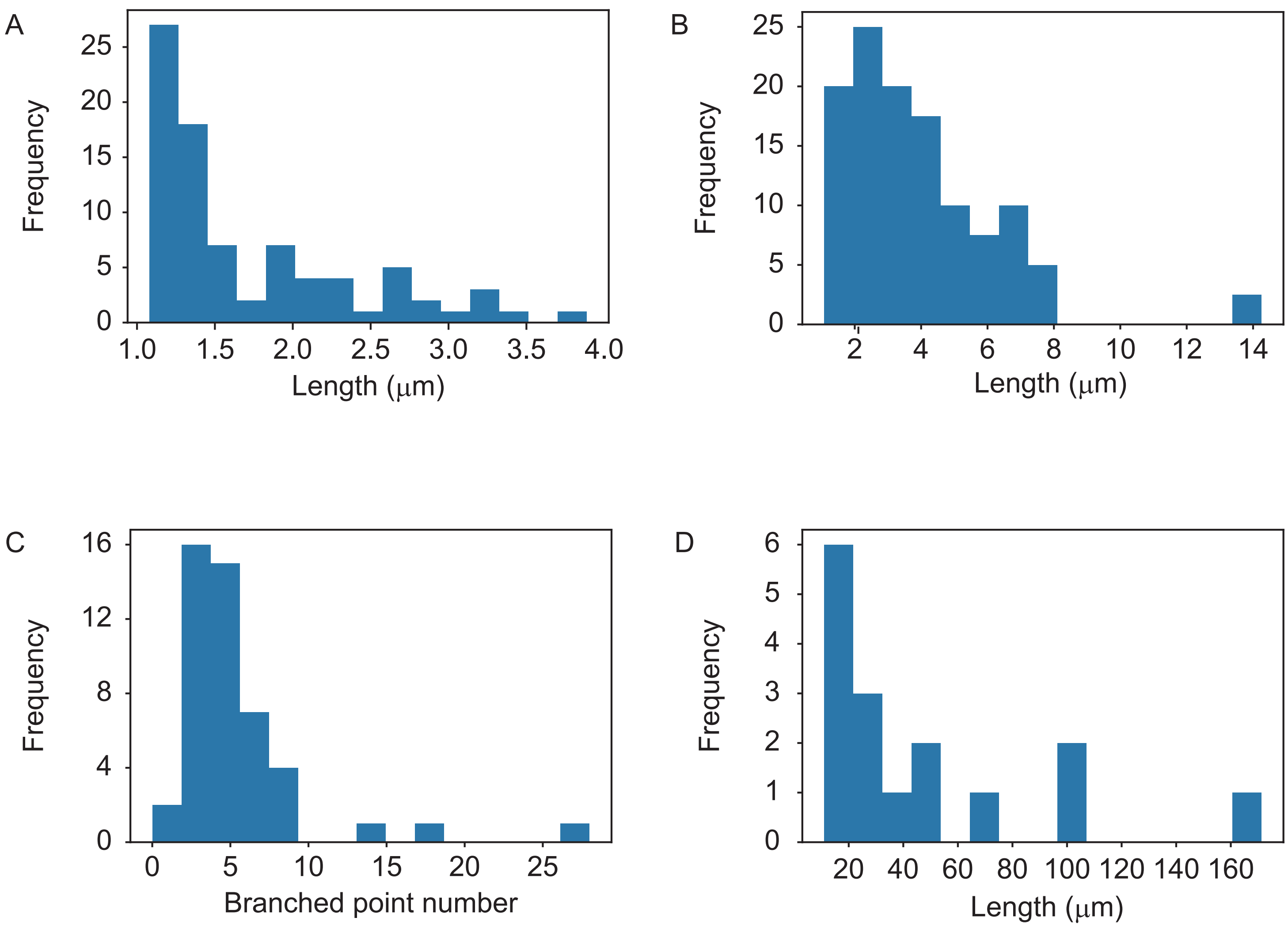


**Supplementary Figure S4. Characteristics of each type of N-ter aggregate**

**(A)** Length of linear fractal-shaped aggregates.

**(B, C)** Length **(B)** and number of branches **(C)** of branched fractal-shaped aggregates.

**(D)** Length of fibril-shaped aggregates.

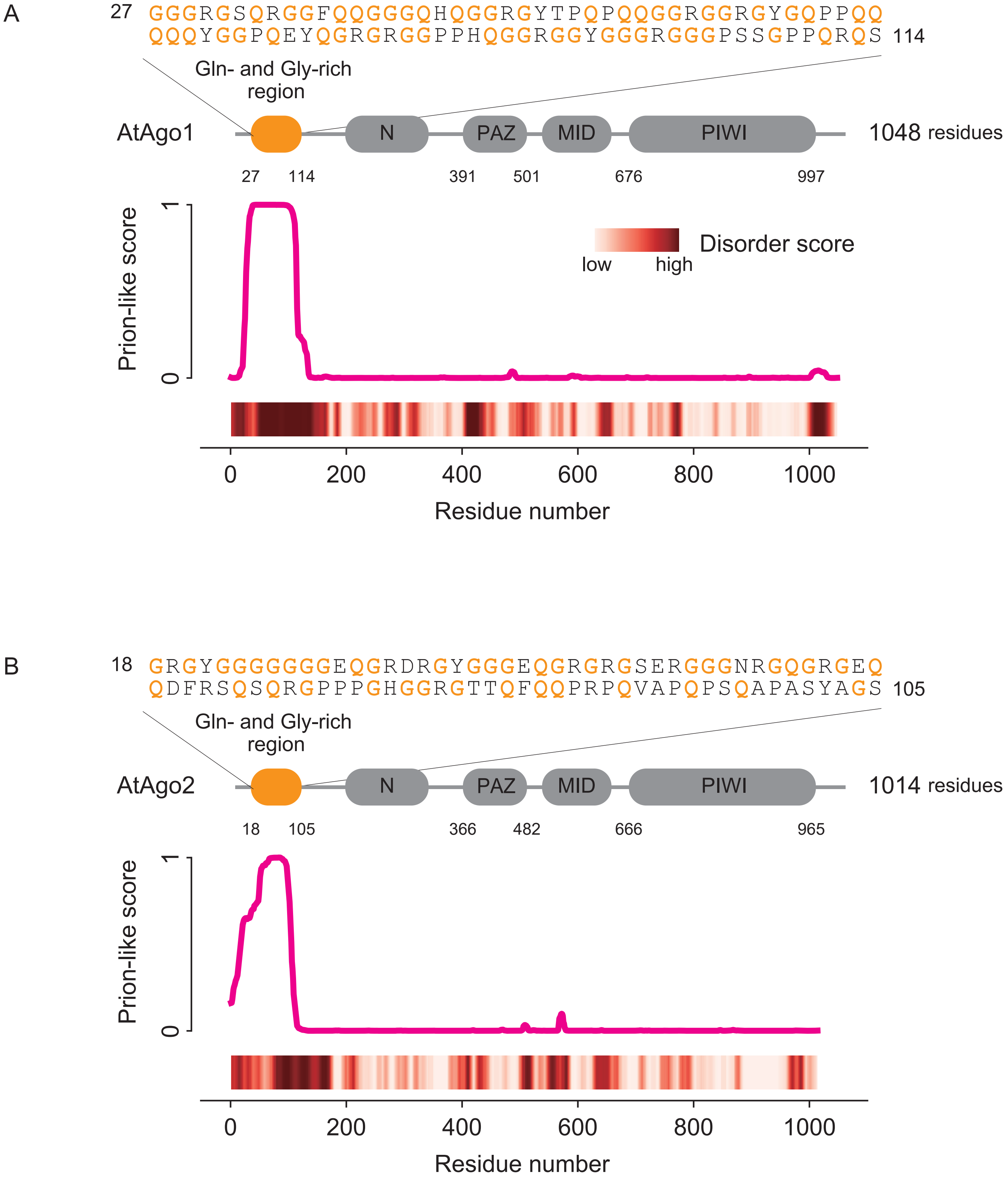


**Supplementary Figure S5. Some plant Argonautes are also predicted to have a NAP domain**

A schematic diagram (top), plot of the prion-like probability sequence predicted by PLAAC (middle) and a heatmap of the degree of disorder predicted by PONDR analysis (bottom) of *Arabidopsis thaliana* Ago1 (**A**, AtAgo1) and Ago2 (**B**, AtAgo2).
